# Engineering of CAR-less lentiviral vectors via ER retention-mediated CAR blockade

**DOI:** 10.64898/2026.06.21.733647

**Authors:** Liya Ma, Jian Wang, Mengwen Huang, Mengjiao Yao, Shuiling Yi, Kaiwen Zhang, Xingyi Ma, Hongxing Jason-Sun

**Affiliations:** School of Biomedical Engineering, School of Science & School of Marine Science and Technology, Harbin Institute of Technology, Shenzhen, Guangdong, China; Shenzhen Celconta Life Science Co., Ltd., Shenzhen, Guangdong, China

## Abstract

Chimeric antigen receptor (CAR)-T cell therapies have transformed the treatment of various tumor types by redirecting and activating T cells against tumor cells. However, CAR-T cell manufacturing approaches remain challenging and limit their widespread use in clinical settings. In vivo CAR-T therapy bypasses ex vivo cell manufacturing and patient preconditioning limitations; however, it faces a significant safety concern as CAR proteins on viral packaging cells are incorporated into budding virions, leading to off-target transduction of tumor cells. Here, we address this risk by developing the CAR-Less ER-Anchor Vector (CLEAN-V) system. By exploiting endoplasmic reticulum (ER) retention, CLEAN-V prevents the CAR protein from trafficking to the cell surface during viral packaging, thereby blocking its incorporation into the viral envelope. CLEAN-V particles exhibit near-complete loss of CAR-mediated tumor cell transduction. Furthermore, CLEAN-V integrates seamlessly into existing third-generation LVV workflows in four- or five-plasmid formats and generates CAR-T cells with preserved phenotypic and functional integrity. These results establish CLEAN-V as a robust platform for developing safe, targeted lentiviral vectors for in vivo CAR-T therapy.

## Introduction

Chimeric antigen receptor (CAR) T cell therapy has recently emerged as one of the most promising alternatives for oncotherapy, especially relapsed/ refractory multiple myeloma (R/R MM)^1–4^. Despite its early promising results, the global adoption of CAR-T therapy remains limited, largely due to complex ex vivo manufacturing processes and the need for patient preconditioning, both of which increase costs and limit patient compliance. Alternatively, in vivo CAR-T cell manufacturing has gained significant attention, as, instead of engineering the CAR-T cells externally, the lentiviral vector (LVV) is delivered directly to the patient ^5–7^. In vivo T-cell engineering occurs within the body, bypassing the intricate ex vivo manufacturing process and the need for preconditioning, a conventional method for creating space that risks depleting immune cells and inducing severe cytopenia ^8,9^. Further, as the T-cells are engineered in situ, they benefit from an optimal physiological environment for survival and expansion, reducing cellular damage often caused by ex vivo expansion and freeze-thaw cycles.

Several CAR-T therapy studies provided evidence of the effectiveness of the in vivo manufacturing approach in circumventing the above limitations. An et al. recently reported a phase 1 study of the evaluation and tolerability of ESO-T01, a nanobody-directed, immune-shielded lentiviral vector (LVV) encoding a humanized anti-B cell maturation antigen (BCMA) CAR for adults with R/R MM, bypassing ex vivo manufacturing or lymphodepleting chemotherapy^10^. Such breakthroughs effectively establish in vivo manufacturing as the next step in CAR-T therapy. However, despite such groundwork, in vivo CAR-T therapies still face several challenges limiting their clinical application, including viral-induced immune toxicities and potential off-target transduction ^11,12^. Currently, the prevailing strategy for engineering targeted lentiviral vectors (tLVVs) involves mutating the Vesicular Stomatitis Virus Glycoprotein (VSV-G) to abolish its native tropism while maintaining its fusogenic activity ^13,14^. In this case, to achieve specificity towards T cells, a single-chain variable fragment (scFv) against CD3, CD7, or other lineage-specific markers is incorporated into the viral envelope ^15–17^. Nonetheless, this strategy presents a significant limitation with regard to the unintentional transduction of tumor cells. This likely occurs because the CAR molecule, co-expressed during the viral packaging process, is incorporated into the viral envelope during budding. This ‘surface CAR’ may then mediate binding to tumor antigens, leading to off-target delivery.

To overcome this obstacle, the CAR protein must be excluded from the packaging cell surface. However, traditional methods fall short as genomic editing permanently disrupts CAR expression while mRNA knockdown strategies often interfere with the viral genomic RNA, resulting in compromised viral titers ^18–21^. Consequently, the direct modulation of the CAR protein itself emerges as the most viable strategy to suppress its surface presence during vector production. In this study, we conducted a proof-of-concept investigation into the application of PEBL to generate CAR-less LVVs, which we term CAR-Less ER-Anchor Vector (CLEAN-V). Protein expression blocker (PEBL) provides an efficient strategy to prevent membrane proteins from trafficking to the cell membrane, resulting in the elimination of their surface expression ^22–24^. Our data demonstrate that PEBL efficiently blocks CAR from trafficking to the packaging cell surface, thereby enabling the generation of CAR-less LVVs. Furthermore, this PEBL platform can be seamlessly integrated into conventional viral packaging workflows using either four-plasmid or five-plasmid transfection systems. Overall, we have developed a novel platform for CLEAN-V production that holds significant promise for in vivo CAR-T cell therapy.

## Results

### CAR-mediated binding enables LVV transduction of tumor cells

During the budding process, lentiviruses acquire their outer envelope from the host cell membrane, resulting in the passive co-packaging of endogenous producer-cell surface proteins onto the viral coat. Using CD19 CAR, we first confirmed the expression of CAR on the packaging cell surface during virus production (Figure 1A). Then, using Enzyme-Linked Immunosorbent Assay (ELISA), we detected the CAR protein within three previously studied CAR LVVs ^25,26^. As expected, CAR proteins were readily detected in all three viral broths. VSV-G can be engineered to ablate low-density lipoprotein receptor (LDLR) binding while maintaining its fusogenic activity, a strategy widely used in targeted lentiviral vector design ^13^. To evaluate the functional impact of CAR proteins incorporated into the viral envelope, we generated anti-CD19 CAR lentiviruses pseudotyped with either wild-type or mutated VSV-G (K47Q, R354Q). We then compared their transduction efficiencies in CD19-negative HEK293T cells versus CD19-positive Raji cells. While the VSV-G mutant (VSVGmut) LVVs exhibit almost complete loss of transduction in HEK293T cells, the anti-CD19 CAR LVVs retained significant transduction capability in Raji cells (Figure 1C). To confirm that this entry was driven specifically by the CAR-target interaction, we performed a competitive protein blocking assay. Strikingly, soluble CD19 protein almost completely abolished transduction of Raji cells by the VSVGmut virus (Figure 1D). Collectively, these data demonstrate that CAR molecules incorporated onto the virion surface during packaging can actively mediate target cell entry for VSVGmut-pseudotyped viruses.

**Figure 1.**
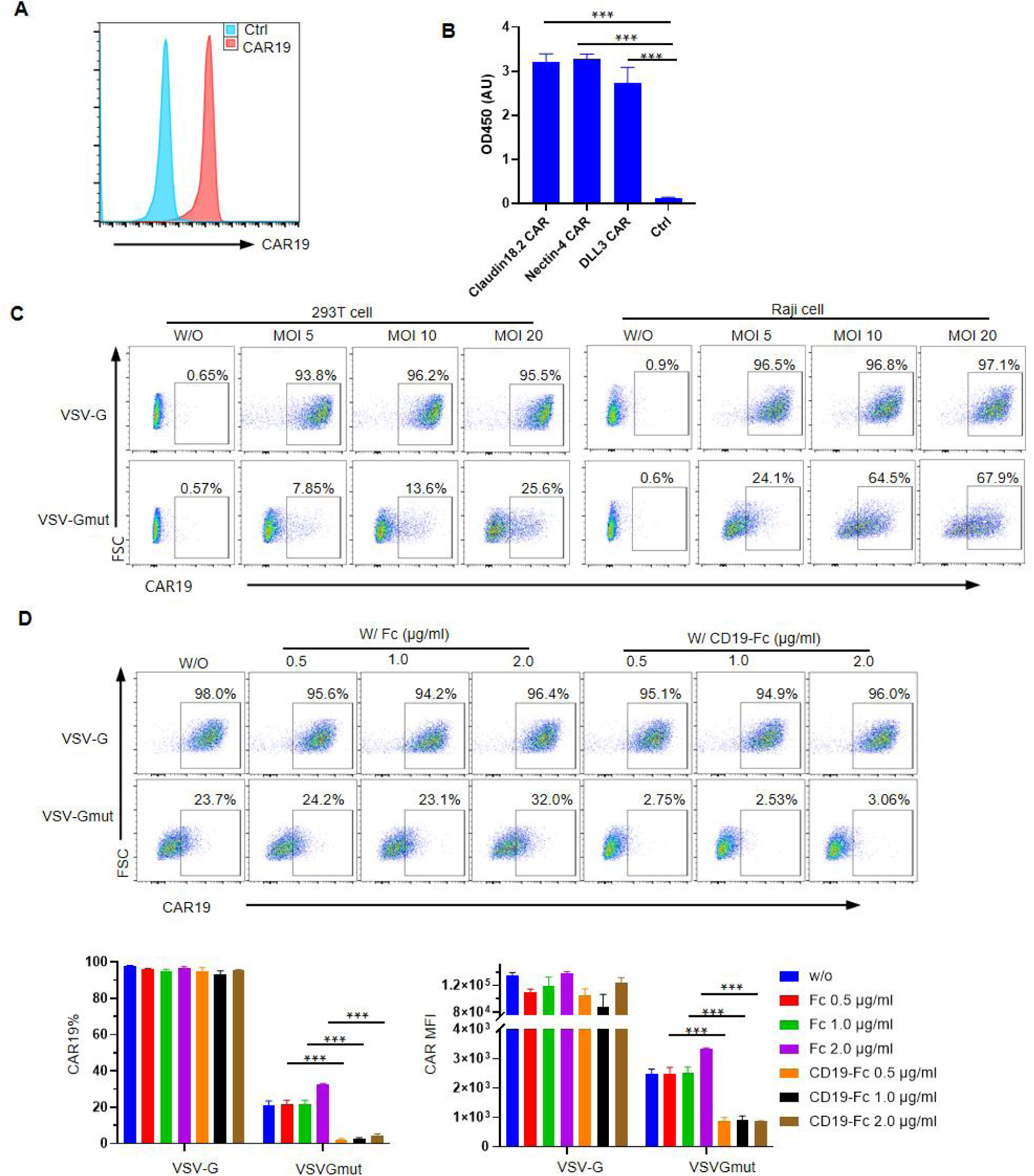
CAR protein on lentivirus vector (LVV) surface can mediate transduction of CAR antigen expressing cells. (A) CAR protein is detected on the surface of HEK293T cells during LVV packaging process. (B) CAR protein is detected in CAR LVV broth but not in non-CAR LVV via ELISA. (C) Transduction of HEK293T cells (CD19-negative) and Raji (CD19-positive) with CAR19 VSV-G and VSVGmut LVVs. (D) CAR-CD19 blockade experiment. Tumor cells were pretreated with CD19-Fc or a control Fc protein for 20 minutes prior to LVV transduction; three days post-transduction, CAR expression was assessed via flow cytometry. All quantitative data are presented as mean ± SD from three independent experimental replicates (n = 3). *** p < 0.001

### Blockade of CAR trafficking using protein expression blocker

Lentiviruses are enveloped RNA viruses. To eliminate CAR display on the producer cell, we sought to manipulate CAR expression at the protein level to ensure the viral genomic RNA undisturbed. To achieve this, we leveraged the endoplasmic reticulum (ER) retention mechanism, which offers an effective method for modulating cell surface protein presentation (Figure 2A). As a proof-of-concept study, an FMC63 scFv-based CD19 targeting CAR was selected as the target construct. To capture the CAR protein, we evaluated two distinct bait proteins—the CD19 protein and an anti-FMC63 antibody derived scFv—each tested in combination with two different ER retention signals (Figure 2B)^23^. To evaluate the system, the PEBL constructs and CAR-encoding plasmids were co-transfected into HEK293T cells, and subsequent CAR surface levels were assessed. As demonstrated in Figure 2C, both scFv-based PEBLs mediated robust CAR surface downregulation, while CD19 protein-based PEBLs showed limited blockade efficiency, suggesting scFv-based bait architecture is superior in capturing and sequestering the intracellular CAR protein. As most of the viral manufacturing workflow spans a 48-hour window, we evaluated the durability of CAR blockade. The surface CAR downregulation was maintained for at least 72 hours, and the PEBL-A signal exhibited superior, long-lasting blockade activity compared to PEBL-B (Figure 2D). To investigate how the bait-to-target protein ratio affects blockade efficiency, we evaluated different transfection plasmid ratios. As expected, we observed a dose-dependent increase in CAR blockade efficiency with increasing amounts of the PEBL plasmid (Figure 2E). Taken together, our data indicate that PEBL can effectively blockade CAR protein trafficking to the cell surface.

**Figure 2.**
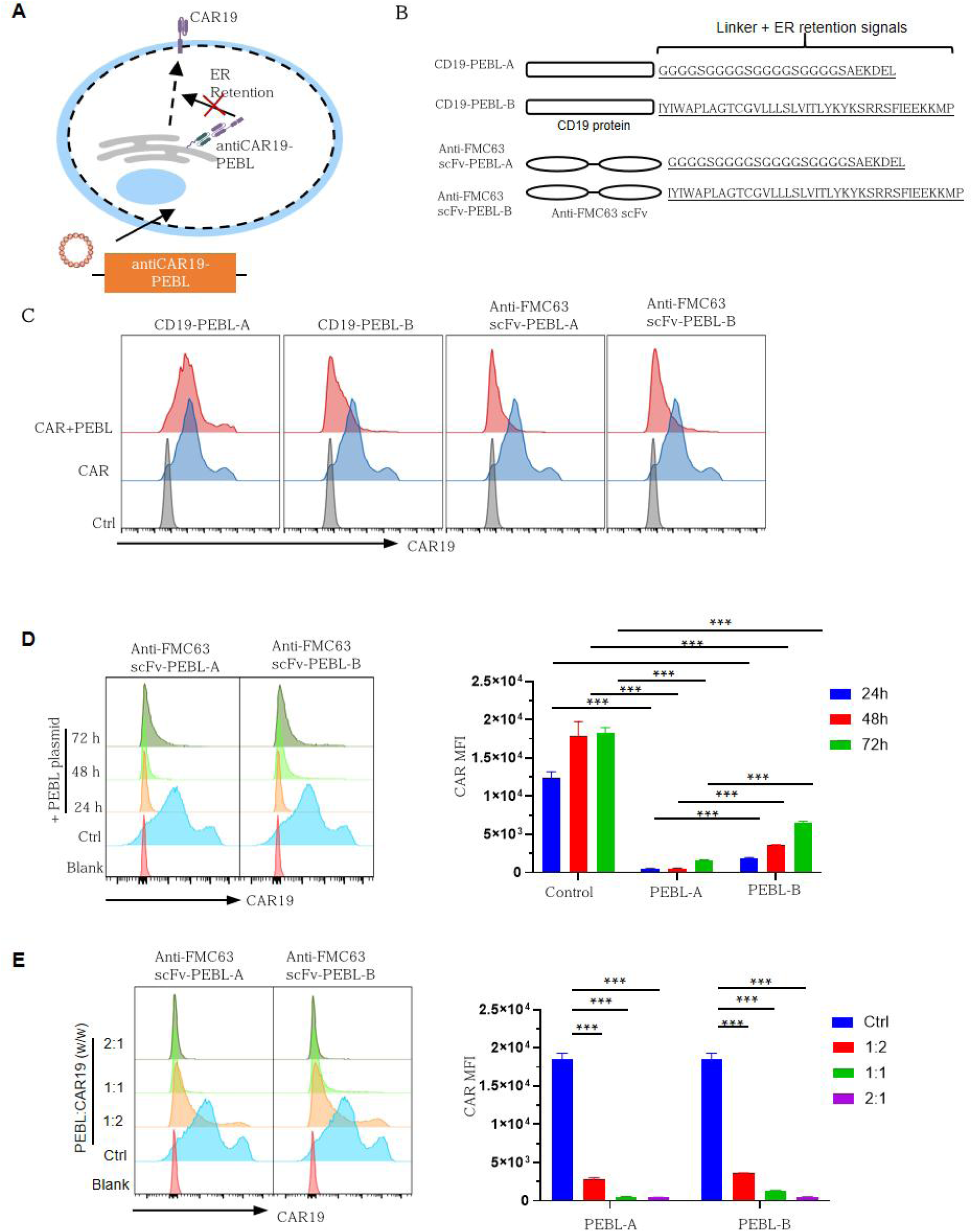
Validation of protein expression blocker (PEBL) mediated CAR blockade. (A) Schematic illustration of PEBL mediated CAR blockade. (B) Schematic of four tested PEBL constructs. (C) Evaluation of PEBL-mediated CAR blockade efficiency. HEK293T cells were co-transfected with a CAR encoding plasmid and a PEBL encoding plasmid at 1:1 mass ratio, followed by flow cytometry analysis of CAR surface expression 48 hours post-transfection. (D) CAR blockade kinetics. To determine CAR blockade kinetics, HEK293T cells were co-transfected with CAR and PEBL plasmids. Surface CAR expression was subsequently analyzed at 24, 48, and 72 hours post-transfection. (E) CAR blockade efficiency is PEBL dose-dependent. HEK293T cells were co-transfected with varying mass ratios of CAR and PEBL plasmids, and surface CAR expression was analyzed 48 hours post-transfection. All quantitative data are presented as mean ± SD from three independent experimental replicates (n = 3). *** p < 0.001

### Incorporation of PEBL into the third-generation lentiviral plasmids

Third-generation lentiviral packaging system consists of four plasmids: Gag/Pol, Rev, VSV-G, and a Transfer vector ^27^. To integrate the PEBL gene into the viral packaging system, several strategies were tested. We first evaluated bicistronic T2A linker to co-express PEBL and VSV-G from a single vector in two alternative orientations. While both configurations mediated surface CAR blockades, the upstream position of PEBL (PEBL-2A-VSVG) demonstrated significantly more robust and consistent blockade efficiency (Figure 3A). We next extended this bicistronic strategy to the Gag/Pol and Rev plasmids. Unfortunately, the modified Gag/Pol construct failed to show any blockade activity, and the REV-P2A-PEBL plasmid yielded only moderate efficiency (Figure 3B). We hypothesized that PEBL transcription might be influenced by adjacent viral genes. To overcome this limitation, we introduced an additional EF1α promoter to drive PEBL expression. Although the Gag/Pol-EF1a-PEBL construct still failed to display any blockade, the blockade efficiency of REV-EF1a-PEBL was significantly improved (Figure 3C). On the other hand, we also tested integrating the PEBL gene into the host genome of the HEK293 cells. HEK293T cells were transduced with a LVV encoding PEBL, and two stable clones were later successfully isolated and verified (Figure 3D). However, subsequent CAR blockade assay yielded very limited CAR downregulation, suggesting insufficient PEBL expression (Figure 3E). Taken together, among the various engineering strategies tested, the PEBL-2A-VSVG and REV-EF1a-PEBL demonstrate effective CAR trafficking blockade.

**Figure 3.**
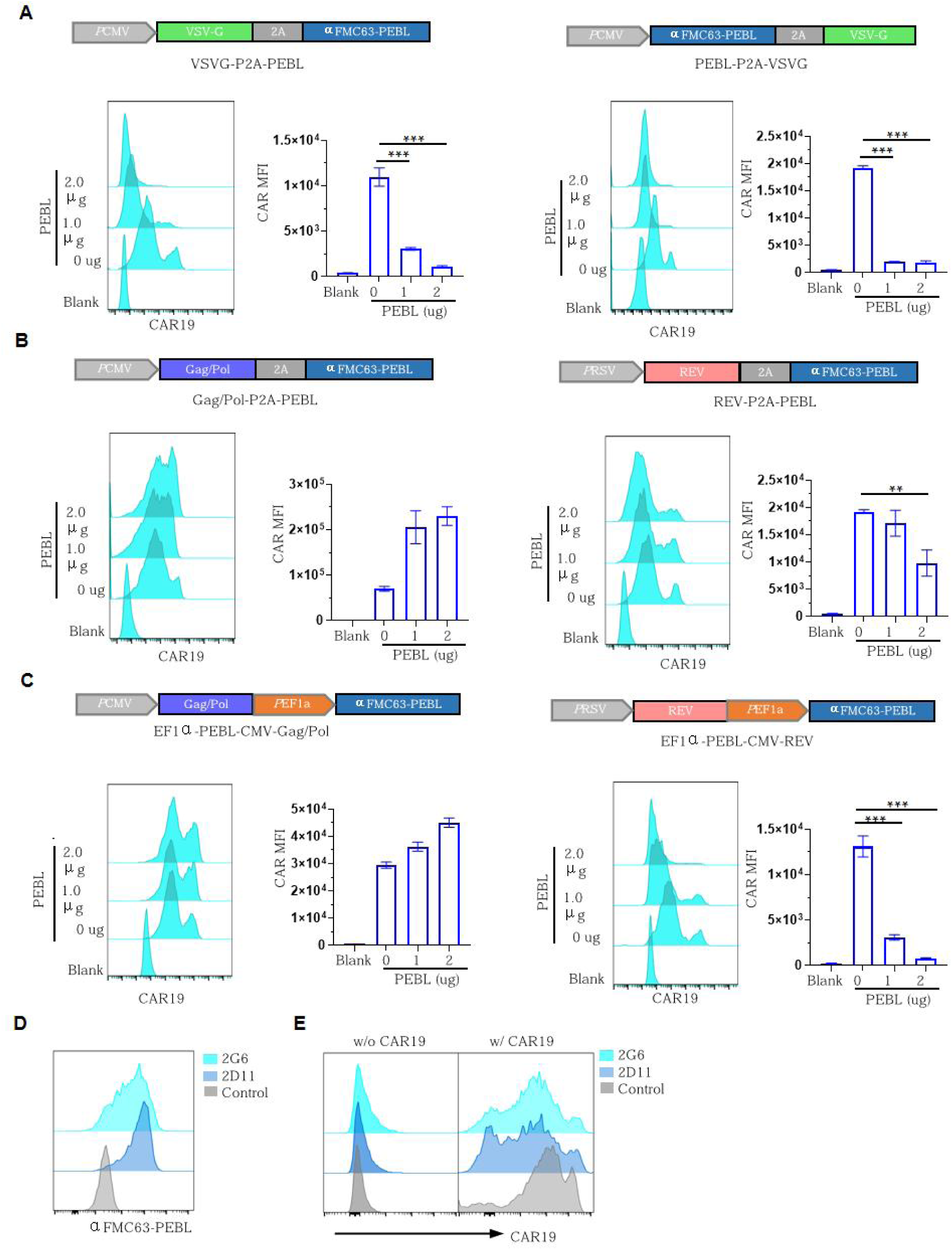
Integration of PEBL into the third-generation LVV system. (A) Integration of PEBL into the VSV-G envelope plasmid. Flow cytometry evaluation of CAR blockade efficiency following transfection with 2A peptide-linked VSV-G and PEBL in two alternate orientations.(B) Flow cytometry analysis of CAR expression following PEBL integration into Gag/Pol and Rev packaging plasmids using a 2A self-cleaving peptide system.(C) Flow cytometry analysis of CAR expression following PEBL integration into Gag/Pol and Rev packaging plasmids using a dual-promoter system.(D) Flow cytometry validation of stable PEBL expression in clonal HEK293T lines (clones 2G6 and 2G11) generated via lentiviral vector (LVV) transduction and subsequent limiting dilution cloning.(E) Evaluation of CAR expression 24 hours post-transfection of the CAR-encoding plasmid into stable PEBL-expressing clones 2G6 and 2G11. All quantitative data are presented as mean ± SD from three independent experimental replicates (n = 3). ** p<0.01; *** p<0.001.

### Production of CLEAN-V using a four-plasmid system

Following plasmid optimization, we next sought to evaluate the efficacy of the PEBL system within a viral production setting. The PEBL-engineered VSV-G and Rev plasmids were transfected into HEK293T cells alongside a CAR-expressing transfer plasmid and a Gag/Pol helper plasmid. The newly engineered vectors were tested either individually or in combination (Figure 4A). While PEBL-VSVG and REV-PEBL each mediated moderate level of CAR blockade, the combination of them achieved the highest blockade efficiency. This is consistent with our finding that the CAR blockade efficiency is PEBL dose-dependent (Figure 4B). Later, lentivirus was successfully generated using the VSVG/REV-PEBL plasmid combination, and CAR protein in the viral broth was quantified using ELISA. Compared to the control group lacking PEBL, virus produced with the PEBL plasmids exhibited significantly lower level of CAR protein (Figure 4C). The vector generated with the PEBL system was designated CAR-Less ER-Anchor Vector (CLEAN-V). Similarly, evaluating CLEAN-V produced with the VSVGmut/REV-PEBL plasmid combination also demonstrated effective CAR blockade (Figure 4D) and residual CAR protein was nearly undetectable in the final viral product (Figure 4E). On the other hand, CLEAN-V titers were reduced by approximately 50% compared to the control virus lacking PEBL (Figure 4F). Finally, we tested the CAR-mediated tumor cell transduction. Compared to the control LVVs, the tumor cell transduction was almost completely abolished with CLEAN-V, suggesting the effectiveness of PEBL system (Figure 4G). Taken together, utilizing four-plasmid system, we successfully generated CLEAN-V, which effectively prevents unintended CAR-mediated transduction of tumor cells.

**Figure 4.**
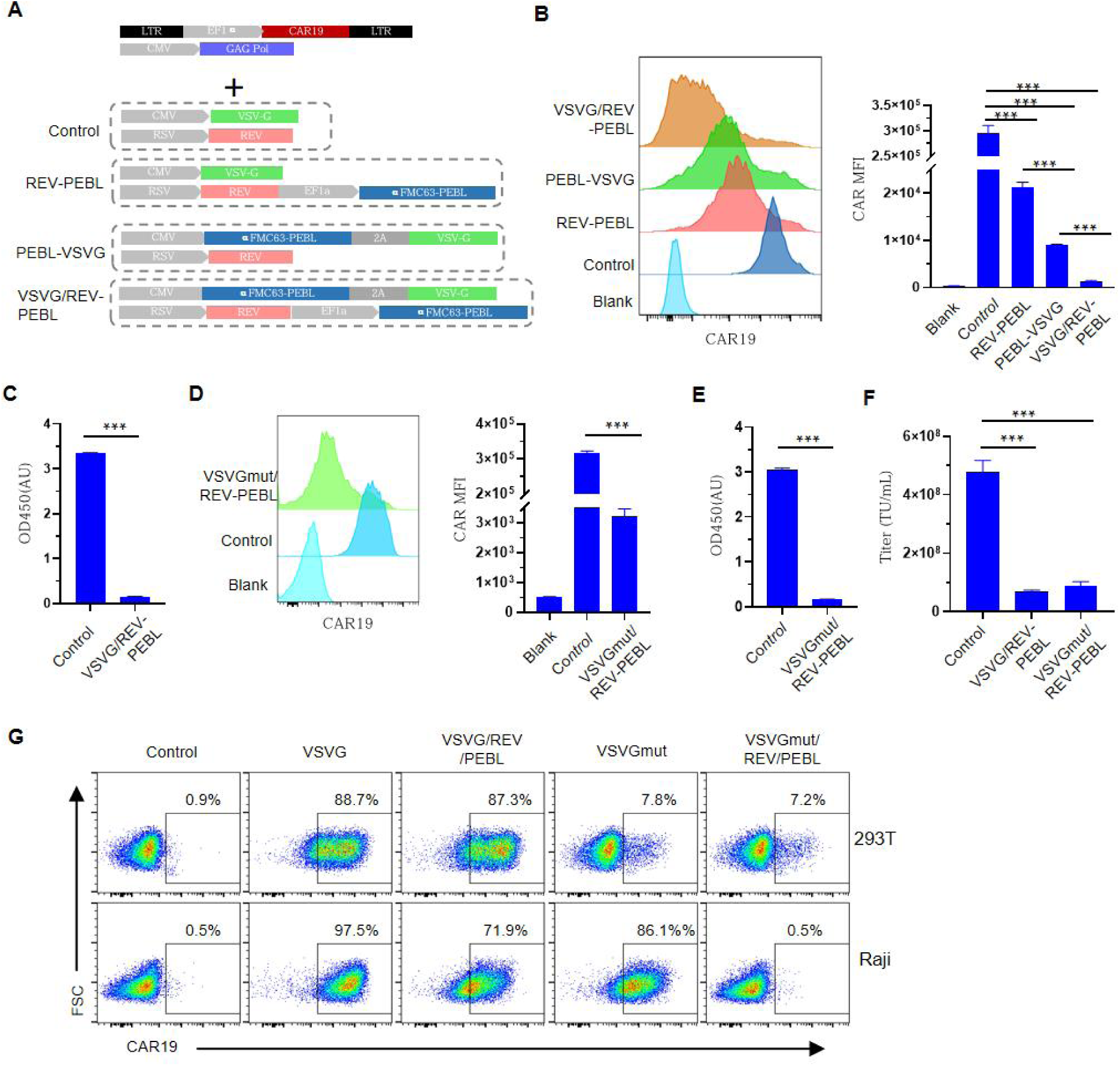
Evaluation of four-plasmid system for CLEAN-V production. (A) Schematic illustration of the plasmid combinations utilized for CLEAN-V production. (B) Evaluation of CAR blockade efficiency across different plasmid combinations during viral packaging. (C) ELISA detection of soluble CAR protein levels within harvested viral particles. (D, E) Production of CLEAN-V using VSV-Gmut plasmid; The CAR expression on HEK293T was determined by flow cytometry (D) and CAR protein in final viral product was determined by ELISA (E). (F) Comparison of viral titers obtained from conventional LVVs versus the CLEAN-V system. (G) Evaluation of the tumor cell transduction of conventional LVVs and CLEAN-Vs. All quantitative data are presented as mean ± SD from three independent experimental replicates (n = 3). *** p < 0.001

### Production of CLEAN-V using a five-plasmid system

Although the four-plasmid system can produce CLEAN-V effectively, it requires custom engineering of the VSV-G and Rev plasmids for each distinct CAR candidate. To establish a more universal platform, we investigated whether introducing a fifth plasmid encoding CAR-PEBL could bypass this limitation, thereby keeping the standard VSV-G and REV packaging plasmids invariant. To test this approach, we constructed a dedicated plasmid carrying the CAR-PEBL gene (Figure 5A) and confirmed that it effectively blocks CAR expression (Figure 5B). Because introducing an additional plasmid could compromise the transfection efficiency of the core packaging components, we titrated the amount of the fifth plasmid during viral production. Increasing the fifth plasmid input progressively enhanced the CAR blockade effect, however, it simultaneously led to a dose-dependent decrease in viral titers (Figure 5C). To balance blockade efficiency with viral yield, we selected 0.27 µg of the PEBL plasmid for subsequent investigations. Furthermore, we evaluated this strategy within the VSV-Gmut system and confirmed the robustness of the five-plasmid approach (Figure 5E). While CAR blockade efficiency was sacrificed to preserve viral titers, residual CAR protein remained nearly undetectable in the final viral product (Figure 5F). Consequently, in tumor cell transduction assays, CLEAN-V generated via the five-plasmid system completely lost its capacity to transduce tumor cells (Figure 5G). Collectively, our data indicate that introducing a fifth PEBL-encoding plasmid can also be utilized for CLEAN-V production, although the plasmid quantity could influence viral titers.

**Figure 5.**
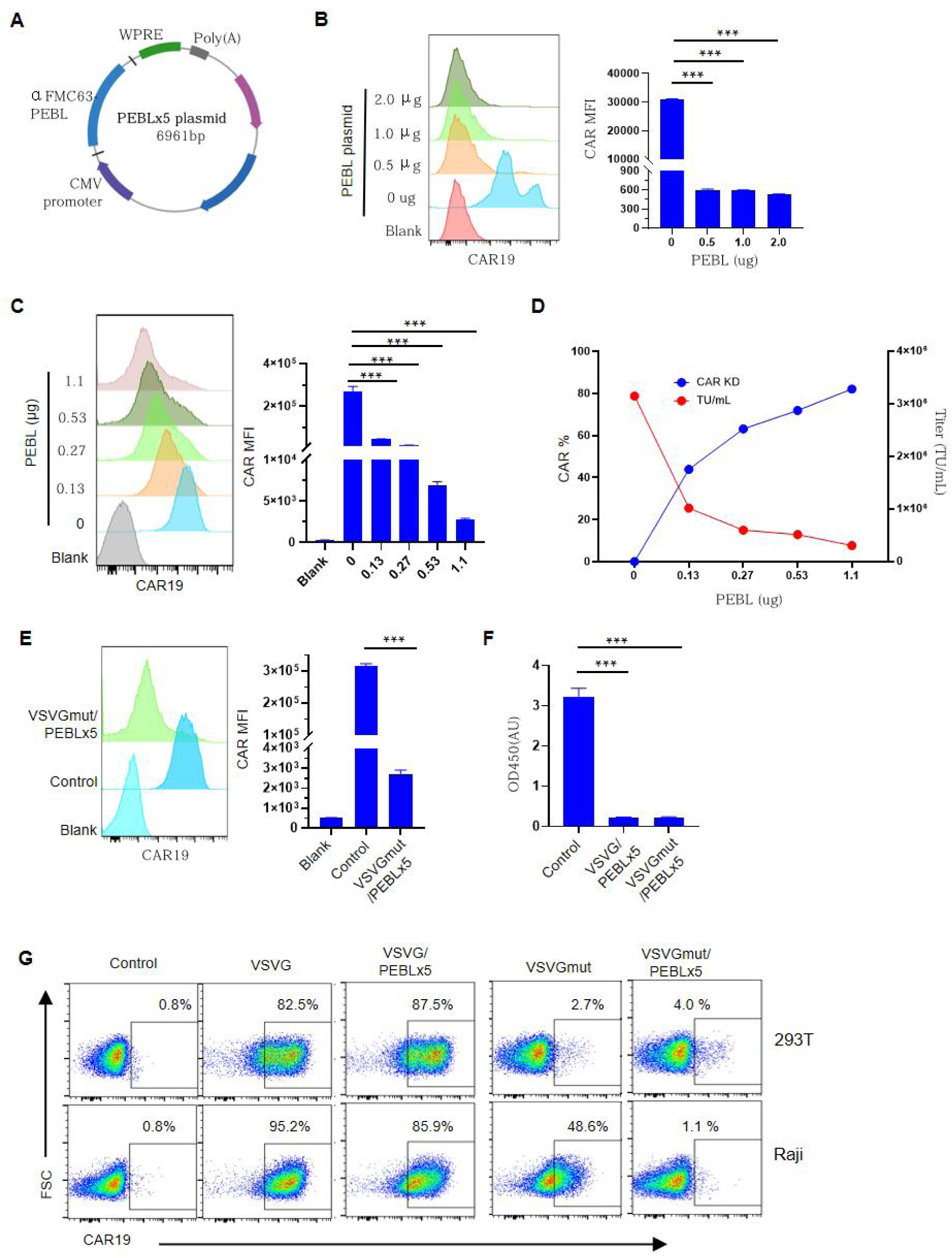
Evaluation of five-plasmid system for CLEAN-V production. (A) Plasmid map of the fifth PEBL (PEBLx5) expression vector. (B) Assessment of PEBLx5-mediated CAR blockade efficiency. (C, D) Titration of the PEBLx5 plasmid for optimal CAR blockade and viral titer. Different amounts of PEBLx5 were co-transfected with four conventional viral plasmids into packaging cells. After 48 hours, CAR expression on HEK293 cells (C) and viral titers (D) were determined. (E) CLEAN-V was produced via a five-plasmid system utilizing VSV-Gmut plasmid, with CAR expression evaluated at the time of viral harvest. (F) CAR protein in final viral product, versus the five-plasmid CLEAN-V system, was determined by ELISA. (G) Evaluation of tumor cell transduction using conventional LVVs versus the five-plasmid produced CLEAN-Vs. All quantitative data are presented as mean ± SD from three independent experimental replicates (n = 3). *** p < 0.001

### Functional characterization of CAR-T cells generated with CLEAN-V

Having demonstrated that CLEAN-V prevents CAR-mediated tumor cell transduction, we next investigated whether it could produce functional CAR-T cells. CLEAN-V was generated using both four-plasmid and five-plasmid systems and subsequently utilized for CAR-T cell manufacturing. Because CAR expression level can influence target recognition and antitumor activity, CAR intensity was carefully evaluated. CAR-T cells generated with CLEAN-V or conventional vectors expressed equivalent levels of CAR protein and displayed similar expression kinetics over the culture period (Figure 6A, 6B). Additionally, the percentage of CAR-positive cells was well maintained during T cell expansion for both CLEAN-V and conventional vectors (Figure 6C). However, despite using identical multiplicities of infection (MOI), CLEAN-V generated via the five-plasmid system demonstrated slightly lower transduction efficiency than that produced with the four-plasmid system or conventional virus (Figure 6A, 6C). Notably, T cells transduced with conventional LVVs and CLEAN-Vs exhibited similar proliferation kinetics (Figure 6D). Analysis of T cell surface markers revealed no significant differences in subset composition between CLEAN-V and conventional vector-transduced T cells (Figure 6E, 6F). To evaluate CAR-T cell function, we conducted an in vitro cytotoxicity assay. As expected, both CLEAN-V and conventional vector-generated CAR-T cells effectively lysed target cells and produced comparable levels of cytokines (Figure 6G, 6H). Collectively, these data indicate that CLEAN-V and conventional lentiviral vectors exert similar effects on T cell phenotype and function.

**Figure 6.**
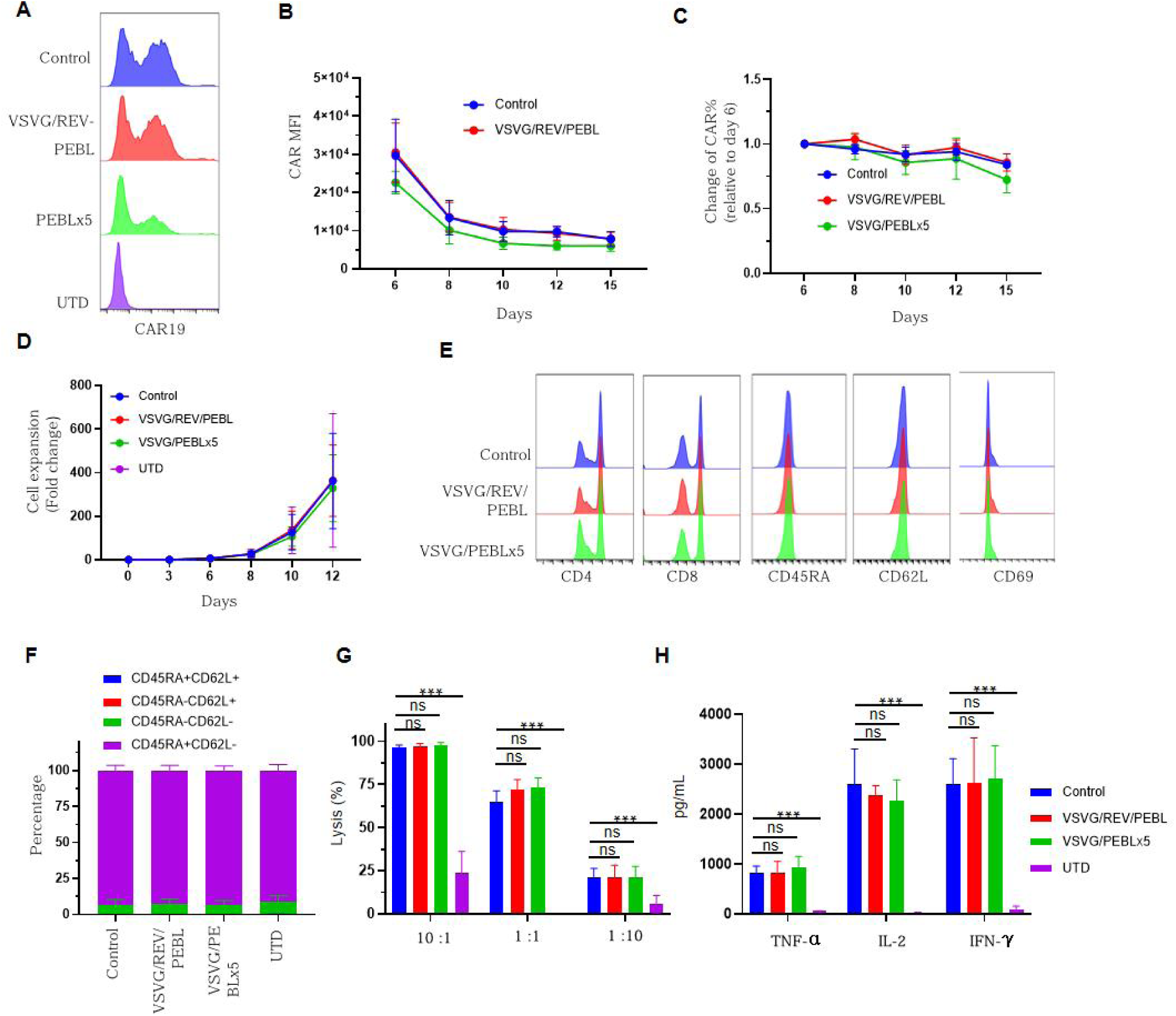
Phenotypic and functional characterization of CAR-T cells engineered with CLEAN-V. Primary T cells were transduced with conventional LVVs or CLEAN-Vs generated using either a four-plasmid or five-plasmid system. (A) Representative CAR expression profiles analyzed on day 6 by flow cytometry. (B) Kinetics of CAR expression intensity and (C) percentage of CAR+ cells over time. (D) T cell growth curves and (E) expression of T cell surface markers. (F) Naïve, effector and memory T cells were analyzed based on CD45RA and CD62L expression. (G–H) In vitro cytotoxicity and cytokine secretion of CAR-T cells engineered with conventional LVVs or CLEAN-Vs. CAR-T cells were co-cultured with Raji-luciferase-expressing target cells at the indicated ratios for 48 hours. (G) Target cell lysis was quantified via luciferase assay, and (H) secreted TNF-α, IL-2, and IFN-γ levels were measured by ELISA. All quantitative data are presented as mean ± SD from three independent experimental replicates (n = 3). *** p < 0.001

## Discussion

As an attractive therapeutic modality, cell therapy has proven its value in the treatment of multiple cancers, particularly hematologic malignancies ^1–4^. However, the extreme complexity of current CAR-T cell manufacturing significantly dampens patient accessibility ^10,12,28^. To tackle these limitations, several strategies have been proposed, including shortening manufacturing times, utilizing RNA- loaded lipid nanoparticle (LNP), and targeted LVVs^7,15,17,29–31^. Among these, targeted LVVs has emerged as a highly promising approach, with early clinical results further supporting its therapeutic potential^15,16^. Nevertheless, a critical concern regarding *in vivo* lentiviral delivery is that the vector may transduce tumor cells via CAR-mediated interaction, leading to the development of CAR-resistant tumor clones ^32^. In this study, we propose a CAR-less LVV generation strategy by leveraging ER retention mechanism ^22–24^. By blocking the CAR protein trafficking at the post-translational level, this approach avoids any disturbance to the viral genome. We have designated this CAR-less vector as “CLEAN-V.” Importantly, CLEAN-V can be generated using either a four- or five-plasmid system, enabling seamless integration into existing vector manufacturing workflows.

*In vivo* generation of CAR-T cells was first investigated using DNA-carrying nanoparticles in 2017 ^33^. Subsequently, Anett Pfeiffer et al. utilized CD8-targeted LVV to generate CAR-T cells in a humanized mouse model ^17^. The recent clinical results reveal that tLVV-mediated *in vivo* CAR-T therapy exhibits promising tolerability and efficacy in treating both tumors and autoimmune diseases ^15,16^. Compared to nanoparticle-mediated transient CAR expression, tLVVs can potentially provide more durable antitumor activity, as their payload integrates into the target cell genome. To achieve precise *in vivo* transduction, the receptor attachment and membrane fusion functions of the viral glycoprotein VSV-G must be separated ^14^. Consequently, several de-targeted VSV-G fusogens have been developed, with target specificity dictated by scFvs against T-cell markers such as CD3, CD4, CD7, and CD8 ^7,13,34^. Because CAR proteins expressed on the viral packaging cell surface can be incorporated into the virion during the budding process, LVVs naturally possess CAR-antigen recognition capability. For ex vivo CAR-T manufacturing, this impact is minimal because VSV-G-mediated transduction remains dominant. However, for tLVV-mediated in vivo CAR-T therapy, this effect is magnified and poses a significant safety concern. Because tLVV transduction relies heavily on its surface-expressed scFvs, tumor cells can be inadvertently transduced via CAR-antigen interactions. This can lead to tumor cell resistance against CAR-T therapy^32^. Therefore, eliminating the CAR protein from LVVs is critical for the development of safe in vivo CAR-T therapies.

Target gene expression can be regulated at various stages of the central dogma, though approach presents distinct challenges for viral engineering. DNA-level modifications using transcription activator-like effector nucleases (TALENs) or clustered regularly interspaced short palindromic repeats associated (CRISPR-Cas) systems permanently alter the target gene sequence, and this irreversibility precludes their use for LVV production ^19–21^. At the transcript level, RNA interference (RNAi)-mediated knockdown can transiently downregulate CAR expression; however, because lentiviruses use RNA as their genome, any perturbation of RNA could compromise final viral titers ^18,35^. Similarly, reversing the CAR expression cassette could segregate the transcribed mRNA from the viral genome; however, this triggers the formation of complementary double-stranded RNA (dsRNA), which interferes with viral packaging ^35^. Ultimately, regulating CAR expression at the protein level emerges as the only viable way for generating CAR-negative vectors.

To optimize CLEAN-V production, we evaluated several strategies. We first attempted to engineer a stable, PEBL-overexpressing HEK293 cell line, thereby avoiding viral plasmid engineering and any modifications to the existing production workflow. However, our results indicate that high blockade efficiency cannot be achieved with this approach, likely due to insufficient PEBL expression. Alternatively, we evaluated four-plasmid and five-plasmid transfection systems. Within the four-plasmid system, the PEBL gene was incorporated into both the VSV-G and Rev plasmids to maximize its expression, while attempts to express PEBL from the Gag/Pol plasmid—via either bicistronic expression or dual-promoter control—failed to induce any CAR blockade. While both systems yield descent viral titers, the four-plasmid system demonstrates superior efficiency. Although neither system achieved a 100% blockade, the CAR protein in the viral broth was nearly undetectable in both; consequently, CAR-mediated tumor cell transduction was almost completely abolished. While the five-plasmid system offers greater operational flexibility, the addition of a fifth plasmid might compromise the transfection efficiency of other plasmids, thereby reducing viral titers. Addressing this limitation may be achieved by reducing the backbone size of the fifth plasmid; the application of a nanoplasmid system, for instance, warrants further investigation One limitation of this platform is that achieving high blockade efficiency requires the selection of optimal CAR-targeting scFvs and ER retention signal sequences. However, any sequence modification to the viral plasmids risks compromising viral production. To circumvent this obstacle, developing a universal CAR-targeting scFv represents an attractive strategy. As most of the CARs utilize a flexible G4S linker to connect heavy and light chains, engineering PEBL that targets this linker could establish a truly universal platform for CLEAN-V production.

In summary, we have conducted a proof-of-concept study demonstrating that the PEBL system can block CAR protein trafficking during LVV production, thereby yielding CAR-less LVVs (CLEAN-V). This platform can seamlessly integrate into existing vector manufacturing workflows. Given the growing enthusiasm for *in vivo* CAR-T cell therapy, our study offers a readily translatable platform for producing safer tLVVs for immunotherapy.

## Materials and Methods

### Cell lines

The human cell lines were obtained from the Cell Culture Center of the Chinese Academy of Medical Sciences (Beijing, China) in the study. HEK293T cells were cultured in Dulbecco’s Modified Eagle Medium (DMEM, Gibco, CA, USA) supplemented with 10% fetal bovine serum (Gibco) at 37 °C in a humidified 5% CO₂ atmosphere. Raji cells were maintained in Roswell Park Memorial Institute 1640 Medium (RPMI-1640, Gibco, CA, USA) supplemented with 10% fetal bovine serum (Gibco) under the same conditions. Primary human T cells were cultured in X-VIVO 15 medium (Lonza, Basel, Switzerland) containing 5% human AB serum (GemCell, CA, USA) and 300 U/mL recombinant human interleukin-2 (IL-2, Thermo Fisher Scientific, MA, USA). All cell lines were routinely tested for mycoplasma contamination by PCR using a commercially available detection kit (Venor™ GeM, Sigma-Aldrich, Germany), and only mycoplasma-free cells were used.

### Plasmid construction

To generate PEBL fusion constructs, the coding sequences of PEBL-A or PEBL-B domains were fused to the 3′end of either the CD19 extracellular domain (signal peptide plus extracellular domain; Uniprot P15391) or the anti-FMC63 antibody fragment. These constructs were subcloned into the pcDNA3.4 vector or the pHBLV lentiviral vector (Hanbio Tech, Shanghai, China). For lentivirus packaging, PEBL constructs were also subcloned into helper plasmids GAG/Pol (Addgene #14887), pRSV-REV (Addgene #12253), and pMD2.G (Addgene #12259), which contained an internal ribosome entry site or a dual EF1α promoter. The VSV-G K47Q and R354Q mutation was introduced into either pMD2.G or PEBL-VSVG plasmid by site-directed mutagenesis using PCR.

### CAR constructs and lentivirus packaging

A chimeric antigen receptor (CAR) construct comprising an anti-CD19 single-chain variable fragment (scFv, FMC63), a CD8α hinge and transmembrane domain, and the cytoplasmic domains of 4-1BB and CD3ζ was inserted into the pHBLV lentiviral vector. For lentiviral vector (LVV) production, the CAR plasmid was mixed with the PEBL construct or with the helper plasmids GAG/Pol, pRSV-REV, and pMD2.G. The DNA mixture was transfected into 293T cells using Lipofectamine 3000 (Invitrogen, Life Technologies, Grand Island, NY, USA). Cells were incubated at 37 °C in 5% CO₂ for 48 h. The supernatant containing viral particles was then harvested, filtered through a 0.45-μm filter, and concentrated using a 100-kDa centrifugal filter unit (Millipore) by centrifugation at 3,500 x g for 45 min at 4 °C.

### DNA transfection and lentiviral transduction

Transfection of HEK293T cells was carried out using Lipofectamine 3000 following the manufacturer’s instructions. For lentiviral transduction, HEK293T and Raji cells were exposed to lentivirus at a multiplicity of infection (MOI) ranging from 5 to 20 in serum-free medium supplemented with CD19-Fc protein (Acro, Beijing, China). After 6 hours, the transduction medium was replaced with fresh medium containing 10% FBS. At 48 hours post-transduction, CAR expression on HEK293T and Raji cells was assessed by flow cytometry using an APC-conjugated anti-G4S linker antibody (B2H1, Hycells, Shanghai, China).

### Generation of CAR-T Cells

Peripheral blood mononuclear cells (PBMCs) were obtained from consenting volunteers through AoNeng Biotechnology (Shanghai, China), adhering to regulatory and institutional guidelines. T cells were isolated from PBMCs and activated using CD3/CD28 Dynabeads (Thermo Fisher Scientific). Activated T cells were then transduced with lentiviral supernatants containing the CAR constructs. After transduction, cells were cultured for an additional two days, after which the CD3/CD28 Dynabeads were removed. CAR-T cells were harvested between days 10 and 13 post-initial activation and used for subsequent cytotoxicity assays.

### T cell cytotoxicity and cytokines release assays

Raji-luc cells (1.0 × 10⁴ cells in 50 µL of RPMI-1640 medium) were seeded per well in 96-well plates. CAR-T cells were then added at effector-to-target (E:T) ratios of 10:1, 1:1, and 1:10 in a total volume of 50 µL. Medium alone served as a negative control. After 48 hours of co-culture, luciferase signal was measured using the GM one-step Luciferase Reporter Gene Assay Kit (Genomeditech, Shanghai, China). Cytolysis was calculated as: cytolysis (%) = [1 – (luminescence of test wells / luminescence of target-only wells)] × 100.

For cytokine release assay, CAR-T cells and Raji cells were co-cultured in triplicate at an E:T ratio of 1:1. At the endpoint of the cytotoxicity assay, supernatants were collected. Levels of interferon-γ (IFN-γ), tumor necrosis factor-α (TNF-α), and interleukin-2 (IL-2) were quantified using a Bio-Plex Pro Reagent Kit (Bio-Rad Laboratories, Inc., CA, USA).

### Flow cytometry assay

The following antibodies were used for flow cytometry: APC anti-human CD3 (clone OKT3), PerCP-Cy5.5 anti-CD69 (clone FN50), PE anti-His tag (clone J095G46) (all from BioLegend, CA, USA); APC-H7 mouse anti-human CD45RA (clone HI100), PE mouse anti-human CD62L (clone DREG-56), APC-Cy7 mouse anti-human CD8 (clone RPA-T8), PerCP-Cy5.5 mouse anti-human CD4 (clone RPA-T4), and PE mouse anti-human PD-1 (all from BD Biosciences, NJ, USA). CAR expression was detected using APC-conjugated anti-G4S linker antibody (B2H1, Hycells, Shanghai, China). Data was acquired on a Beckman Coulter flow cytometer and analyzed with FlowJo software (LLC, OR, USA).

### Statistical analysis

Data are presented as mean ± standard deviation (SD) unless otherwise stated. All experiments were performed with at least three independent biological replicates. Statistical analyses were performed using GraphPad Prism 10 (GraphPad Software, Inc., La Jolla, CA, USA). Comparisons between two groups were analyzed using a paired, two-sided t-test unless otherwise indicated. P value < 0.05 was considered statistically significant.

## Acknowledgements

This work was supported by Shenzhen Celconta Life Science Co., Ltd.

## Author contributions

H. S. designed and directed the experiments. L. M., M. H., K. Z. and J. W. performed the experiments. H. S., L. M., X. M. and J. W. analyzed the data and wrote the manuscript. H. S. supervised the project. All authors read and approve the final manuscript.

## Declaration of generative AI and AI-assisted technologies in the writing process

During the preparation of this work the authors used DeepSeek and Gemini to improve readability and language. After using these tools, the authors reviewed and edited the content as needed and take full responsibility for the content of the publication.

## Data availability

The data that support the findings of this study are available from the corresponding author upon reasonable request.

